# Autophagy as a mechanism for adaptive prediction-mediated emergence of drug resistance

**DOI:** 10.1101/2021.02.02.429461

**Authors:** Nivedita Nivedita, John D. Aitchison, Nitin S. Baliga

## Abstract

Drug resistance is a major problem in treatment of microbial infections and cancers. There is growing evidence that a transient drug tolerant state may precede and potentiate the emergence of drug resistance. Therefore, understanding the mechanisms leading to tolerance is critical for combating drug resistance and for the development of effective therapeutic strategy. Through laboratory evolution of yeast, we recently demonstrated that adaptive prediction (AP), a strategy employed by organisms to anticipate and prepare for a future stressful environment, can emerge within 100 generations by linking the response triggered by a neutral cue (caffeine) to a mechanism of protection against a lethal agent (5-FOA). Here, we demonstrate that mutations selected across multiple laboratory evolved lines had linked the neutral cue response to core genes of autophagy. Across these evolved lines, conditional activation of autophagy through AP conferred tolerance, and potentiated subsequent selection of mutations in genes specific to overcoming the toxicity of 5-FOA. We propose a model to explain how extensive genome-wide genetic interactions of autophagy facilitates emergence of AP over short evolutionary timescales to potentiate selection of resistance-conferring mutations.

## INTRODUCTION

Antimicrobial resistance has become a major threat to modern medicine with ever adapting strains of microbes and evolution of sub-populations of treatment resistant cells. According to CDC, in the U.S. alone, at least 2.8 million people per year contract an antimicrobial-resistant infection^1^. Drug resistance is also a growing concern in the treatment of cancers. For the development of efficacious treatment regimen that accomplishes complete clearance of the pathogens and tumor cells, it has, therefore, become imperative to determine the evolutionary trajectories and associated mechanisms that enable the selection of rare resistance-conferring mutations. Recent evidence has shown that the selection of resistance conferring mutations is potentiated by a preceding tolerant state that may manifest from drug-induced physiological adaptation or mutations in generalized tolerance networks^2,3^. Here we have investigated how emergence of resistance may be potentiated by adaptive prediction (AP), a phenomenon utilized by all organisms to anticipate and prepare in advance for a future environmental change^4–6^. For example, *E. coli* and *M. tuberculosis* sense neutral cues such as rise in temperature or a drop in oxygen to adaptively predict a hostile host environment^4,5^. In other words, AP may allow an organism to transiently tolerate stressful environments including the lethal effects of drugs, thereby increasing the likelihood of selecting drug-specific resistance mutations.

Previously, we demonstrated that repeated exposure to a neutral cue followed by a sub-lethal dose of a toxin drives the emergence of AP-mediated tolerance over short evolutionary cycles^6^. Specifically, over every 10 generations we subjected *S. cerevisiae* (yeast) to pre-conditioning with a neutral dose of caffeine (environmental cue) followed by a sub-lethal dose of 5-fluoroorotic acid (5-FOA; a lethal drug) (Fig 1a)^6^. 5-FOA is toxic to yeast because upon conversion by Ura3 to 5-fluorouracil (5-FU, a pyrimidine analogue), 5-FU is misincorporated into RNA and DNA in place of uracil. The coupled treatments with caffeine and 5-FOA were therefore interspersed with a period of counter-selection in a growth medium without uracil to weed out 5-FU resistant uracil auxotrophs (e.g., *ura3-* mutants). The cyclic exposure to the novel coupled environment of caffeine followed by 5-FOA, resulted in emergence of AP within 50-150 generations across all evolved lines^6^. Remarkably, across all lines AP was eventually displaced by a population that had higher overall resistance to 5-FOA. AP was also shown to emerge from rewiring of regulatory and signaling networks by caffeine-induced segregation of Ura3 from the cytosol to peroxisomes^6^. However, not all of the engineered evolved lines gained AP through this rewiring mechanism suggesting the existence of additional mechanisms for AP-mediated 5-FOA tolerance. Here, we analyzed selected mutations and differentially expressed genes in evolved lines and discovered that the mechanism of AP across multiple evolved lines could be explained by rewiring of the caffeine response to the autophagy network.

**Figure 1.**
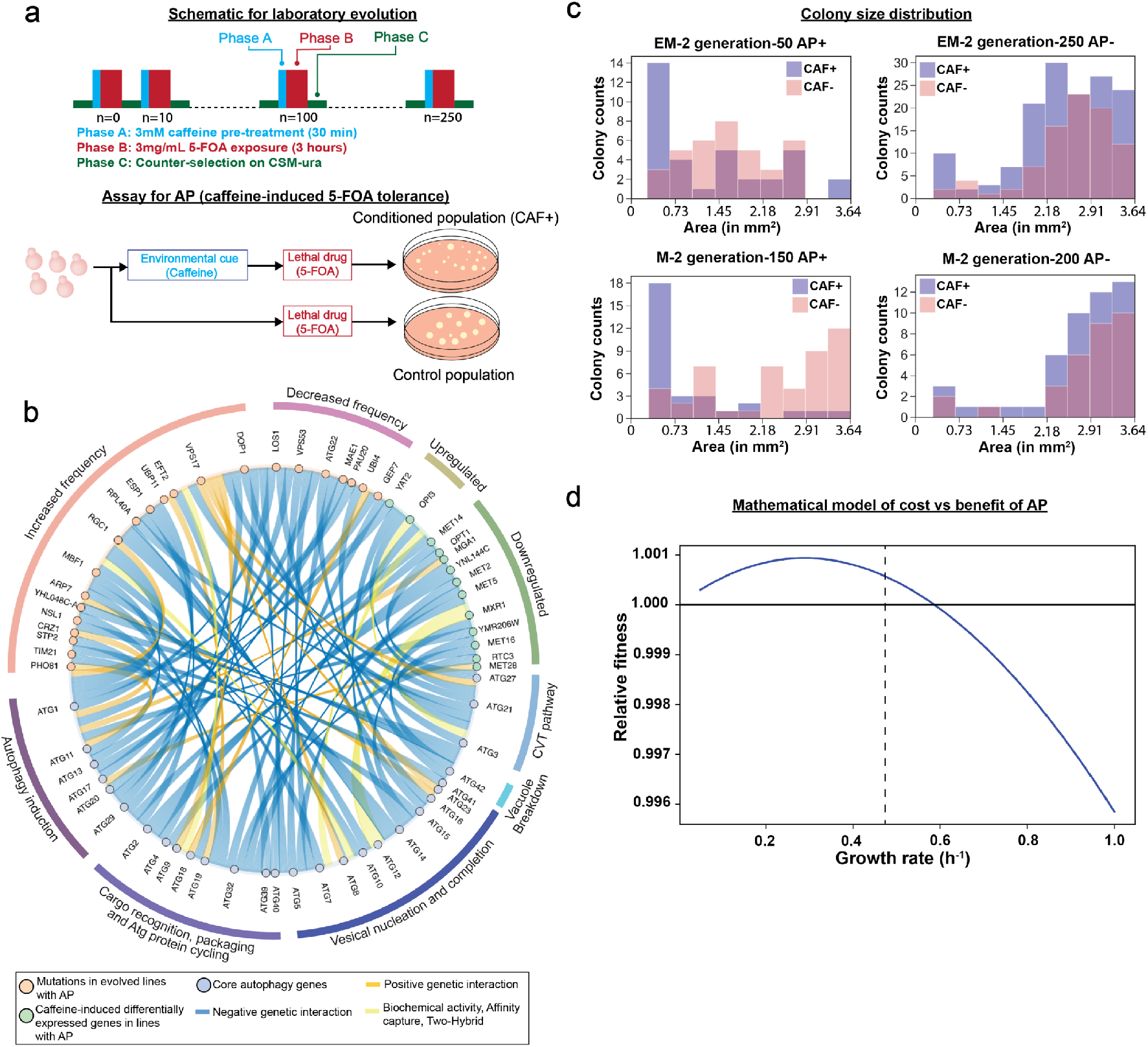
Regulation of emergence of adaptive prediction (AP) during laboratory evolution. (**a**) Experiment design for laboratory evolution of caffeine-induced AP of increased conditional fitness to 5-FOA. (**b**) Genes with increased or decreased mutation frequency or perturbed regulation in evolved lines with AP (relative to generation 0) interact genetically, physically, or biochemically with core autophagy genes. (**c**) Distribution of colony sizes in mutagenized evolved lines of both engineered (EM2) and wild type yeast (M2). Colony size distribution indicates presence of smaller sized colonies in caffeine-treated (CAF+) cultures of evolved lines with AP. Colony sizes were measured using Image J plugin Trainable Weka segmentation and the pixel area was converted to mm^2^ using a scaling factor calculated based on total pixel area of the entire plate. (**d**) Relationship between growth rate and relative fitness inferred by a mathematical model of cost vs benefit of AP. Vertical dashed line indicates growth rate of wild type yeast.

## RESULTS AND DISCUSSION

To determine mechanisms of AP, we analyzed whole genome sequences and RNA-seq data for up to 5 evolved lines across generations over which they gained AP capability (supplementary data from Lomana *et al.^6^*). This analysis discovered mutations in 95 genes (90 genes with ≥20% frequency) and caffeine-induced >2-fold differential expression of 39 genes that may have contributed to the AP phenotype. Using YeastMine^7^ we discovered that 13 differentially expressed genes either regulate or genetically interact with AuTophaGy (ATG) genes (Fig 1b). Notably, 9 out of 11 downregulated genes have negative genetic interactions with ATG genes, suggesting that caffeine-induced downregulation of these genes might have induced autophagy. In parallel, using PROVEAN^8^ to infer consequences of mutations on protein function, we discovered 24 deleterious/non-synonymous and upstream intergenic mutations in 34 genes (including non-synonymous mutation in VPS17 and upstream gene variant in ATG22) that also genetically interact with ATG genes (Fig 1b). Together the mutation and differential gene expression analysis implicated caffeine-induction of autophagy as a plausible mechanism by which multiple evolved lines had gained AP.

Autophagy is an evolutionarily conserved process for recycling amino acids and nutrients from damaged proteins and other cellular macromolecules to support cell survival in stressful environments^9,10^. In yeast, induction of autophagy is associated with decreased growth rate, which manifests in smaller sized colonies in stressful or lethal environments^9,10^. Strikingly, we observed that caffeine pre-treatment resulted in higher numbers of smaller colonies, in generations when the evolved line exhibited AP (Fig 1c). Using the deterministic model for AP developed by Mitchell et al.^11^, we determined that reduction in growth rate in response to a preceding neutral cue could indeed confer higher conditional fitness to a future stressful environment, bolstering caffeine-induced autophagy as a possible mechanism for AP (Fig 1d).

To generate definitive evidence that AP had emerged from rewiring of caffeine response to the autophagy network, we performed the Rosella assay using a dual-color emission biosensor^12^. The Rosella biosensor consists of two tandem fluorescent proteins- a pH sensitive green fluorescent protein (GFP) and a pH stable red fluorescent protein (RFP). This biosensor is targeted to the pH neutral cytosol under normal conditions, wherein both GFP and RFP are active. Upon induction of autophagy, the biosensor is translocated to an acidic vacuole wherein the pH-sensitive GFP is selectively inactivated and only the RFP remains active. Hence, the ratio of red to green fluorescence can be used as a measure of bulk autophagy. Using the Rosella assay, we determined that autophagy was induced upon caffeine treatment across multiple evolved lines in generations exhibiting AP, but not in parental cultures. The longitudinal patterns of changes in caffeine-induced autophagy and conditional fitness to 5-FOA were remarkably correlated (Fig 2a). This observation suggested that AP had emerged through the selection of mutations that mechanistically linked the caffeine-response to autophagy to adaptively predict and increase survivability to 5-FOA. Notably, the loss of AP across the evolved lines was accompanied by correlated reduction in caffeine-induced autophagy and an overall increase in resistance to 5-FOA through the selection of mutations in URA2 and URA6^6^.

**Figure 2.**
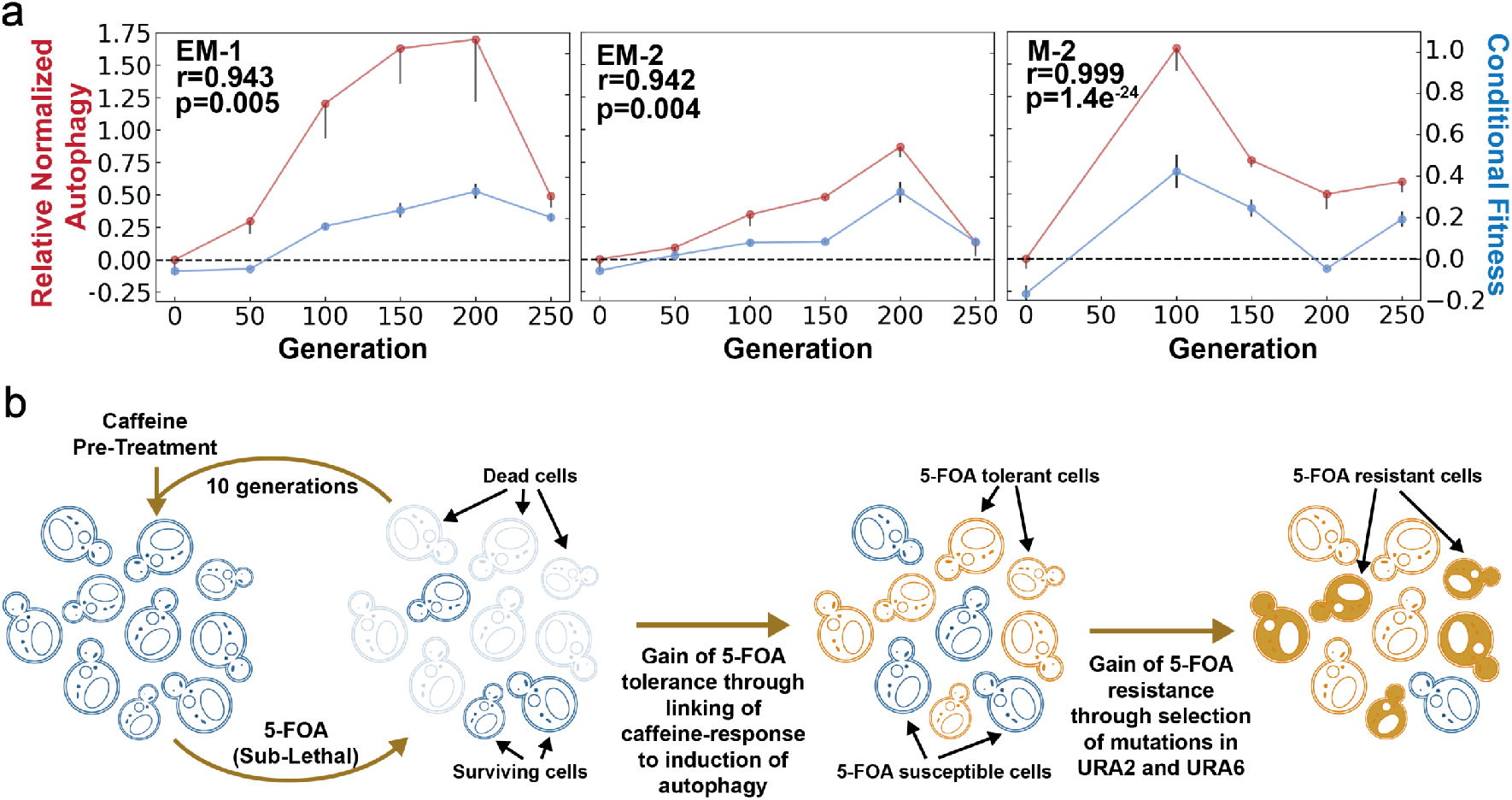
Evaluation of autophagy as a mechanism for adaptive prediction. (**a**) Relative autophagy and conditional fitness in EM and M cell lines. AP emerged within 100-150 generations in all evolved lines. Spearman correlation (r) between conditional fitness and relative autophagy, and associated significance (p-value), is indicated for each evolved line. (**b**) Model for how the rewiring of a neutral cue-induced response (caffeine in this study) to autophagy is a generalized mechanism for generating AP-based tolerance and subsequent resistance to a lethal stress. Rise of a tolerant population due to AP could ultimately potentiate development of resistance through subsequent selection of mutations in specific genes and pathways targeted by the lethal stress (in this example, mutations in Ura2/Ura6 genes of uracil biosynthesis pathway for constitutive resistance to 5-FOA).

Our results demonstrate that mutations selected across at least 3 evolved lines had linked the response to pretreatment with a neutral dose of caffeine to induction of autophagy. While the induction of autophagy may have reduced flux through uracil biosynthesis pathway by generating uridine phosphates^13^, evidence generated in this study has demonstrated that autophagy-mediated reduction in growth rate is also a plausible mechanism of 5-FOA tolerance. Notably, 1,886 of the 6,604 total genes in the yeast genome genetically interact with the 33 genes that are implicated in the core autophagy process, explaining why multiple evolved lines had converged on the same solution for AP-mediated 5-FOA tolerance. We posit that pathogens might exploit rewiring of host-relevant environmental cues to autophagy as a mechanism to rapidly evolve AP-mediated generalized tolerance to diverse lethal agents. The advanced preparedness through conditional induction of autophagy creates a window of opportunity for cells to encounter and select resistance-conferring mutations in genes directly associated with the mechanism of action of a given drug (Fig 2b). This model offers an adaptive evolutionary perspective for the wide association of autophagy with the development of drug resistance^14^, and extends implications of this phenomenon to emergence of resistance in pathogens. Further, the model also proposes that autophagy suppressor drugs could improve efficacy of treatment regimen by blocking a major evolutionary pathway for gain of drug resistance^15^.

## MATERIALS AND METHODS

### Yeast strains, culturing, and conditional fitness measurements

All strains used in this study were derived from the parental BY4741-URA3 strain of *S. cerevisiae*, and reported previously in Lomana et al.^6^ “EM” and “M” refer to lines that originated from UV-mutagenized cultures of the engineered and parental BY4741-URA3 strains mutagenized as described in Lomana et al.^6^ Glycerol stocks for generations 0-250 for each evolved line were revived on agar plates, cultured overnight to log-phase in complete synthetic media (CSM)-uracil with 2% glucose, at 30°C on a shaker. Cell density was determined by hemocytometer counts. Conditional fitness assays were performed as described previously by Lomana *et al.^6^*. In brief, a culture aliquot was subjected to a sub-lethal dose of 5-FOA (3mg/mL) for 3 hours, with and without 30 minutes of pre-conditioning with 3mM caffeine. Relative change in 5-FOA survival (*S*) with vs. without caffeine pre-treatment was used to calculate conditional fitness (CF), where S is calculated using equation 1:

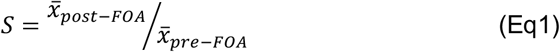

where 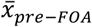 and 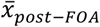 are average CFU counts before and after 5-FOA treatment.

Conditional fitness (CF) is calculated as difference between survival with caffeine pretreatment (Sc) and survival without caffeine pre-treatment or control (*S_o_*).

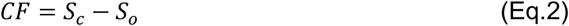

### Autophagy Measurement

Cells from generations 0-250 for each evolved line were transformed with the Rosella biosensor, cultured to log phase, and concurrently evaluated for AP and autophagy. After subtraction of background fluorescence, overall autophagy was calculated as the difference in ratio of average red to green fluorescence, with and without (control) caffeine pretreatment, and plotted in reference to overall autophagy in the parental population. Autophagy difference (A_d_) was calculated as follows:

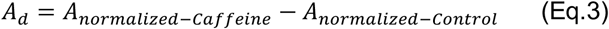

where *A_normalized-Control_* and *A_normalized-Caffeine_* are autophagy measurements normalized relative to the average autophagy in the parental population. Relative normalized autophagy was then determined relative to A_d_ at Generation 0.

## ACKNOWLEDGMENTS

We thank Adrian Lopez Garcia de Lomana and Thurston Herricks for comments and feedback on experimental design. This work was supported by the National Institutes of Health grant 5R01AI141953 to N.S.B and J.D.A. and 5R01AI128215 to N.S.B.

